# Visualizing the superfamily of metallo-β-lactamases through sequence similarity network neighborhood connectivity analysis

**DOI:** 10.1101/2020.04.16.045138

**Authors:** Javier M. González

## Abstract

The superfamily of metallo-β-lactamases (MBL) comprises an ancient group of proteins found in all domains of life, sharing a characteristic αββα fold and a histidine-rich motif for binding of transition metal ions, with the ability to catalyze a variety of hydrolysis and redox reactions. Herein, structural homology and sequence similarity network (SSN) analysis are used to assist the phylogenetic reconstruction of the MBL superfamily, introducing tanglegrams to evaluate structure-function relationships. SSN neighborhood connectivity is applied for spotting protein families within SSN clusters, showing that 98 % of the superfamily remains to be explored experimentally. Further SSN research is suggested in order to determine their topological properties, which will be instrumental for the improvement of automated sequence annotation methods.

---

The metallo-β-lactamase (MBL) superfamily comprises an ancient group of proteins found in all domains of life, sharing a characteristic αββα fold and a histidine-rich motif for binding of transition metal ions. The name was coined after the first superfamily members to be characterized; a group of zinc-dependent hydrolases produced by bacteria resistant to β-lactam antibiotics. These zinc-β-lactamases (ZBLs) hydrolyze the amide bond present in all β-lactams and thus render them ineffective. The first X-ray crystallographic report of a ZBL was that of BcII from *Bacillus cereus* 569/H/9 [1].

Despite its low resolution, the atomic model disclosed the new αββα fold and a single Zn(II) ion bound to a three-histidine motif, resembling the active site of carbonic anhydrases. Thus, BcII and ZBLs in general were believed to use a single Zn(II) ion to activate a water molecule for hydrolysis, paralleling the mechanism by which carbonic anhydrases catalyze carbon dioxide hydration. This hypothesis was soon questioned when the structure of ZBL CcrA from *Bacteroides fragilis* was published, disclosing a bimetallic zinc center, with the second zinc being coordinated to nearby Asp, Cys and His residues [2]. Besides, the second zinc was later found in *B. cereus* ZBL too [3–5], starting a decade-long controversy regarding the role of each zinc ion. Later on, it was found that monometallic ZBLs are rather exceptional and the hydrolysis reaction generally requires two Zn(II) ions [6, 7]. It is important to note that, while ZBLs hydrolyze antibiotics by means of a metal-activated water molecule, most β-lactamases use a conserved serine residue in a completely different protein scaffold. In other words, the majority of β-lactamases are not metallic, and referring to ZBLs and MBLs in general simply as “β-lactamases” should be avoided, particularly when annotating these proteins in public databases. Besides, even though most members of the superfamily are devoid of β-lactamase activity, the acronym MBL has been adopted to annotate most members of the superfamily. The same convention will be followed here to define any protein with at least one characteristic MBL domain, leaving the acronym ZBL to describe metallo-β-lactamases themselves.

A great diversity of proteins evolved in the MBL superfamily by combining catalytic MBL domains and substrate recognition domains in a modular fashion. Besides, subtle changes in the metal coordinating residue networks expand this diversity by enabling the coordination of different transition metals, particularly Zn(II), Mn(II), and Fe(II)/Fe(III) (Figure 1). Early attempts to build a systematic classification of the MBL superfamily were conducted by L. Aravind [8], as some of the very first applications of the PSI-Blast algorithm [9], who showed that many proteins other than ZBLs comprise the characteristic fold and histidine-rich metal-binding motif of MBLs, mapping key residues onto the structure of *B. cereus* ZBL. These observations were updated in 2001 by Daiyasu *et al.*, when additional crystal structures of MBL superfamily members were available [10]. At present, more than a hundred proteins have been shown to contain αββα domains through X-ray crystallography, whereas the InterPro 77.0 database [11] entry IPR001279 for the MBL superfamily includes about half a million members. Indeed, the MBL superfamily has grown astoundingly over the past 30 years, and an integrative revision is long overdue. In this report, the Pfam and Protein Data Bank databases are interrogated in order to obtain an updated picture of the distribution of proteins throughout the MBL superfamily and to get insights into their structure-function relationships. It is shown that the most widespread MBLs are those acting on phosphoesters, such as nucleic acids, nucleotides, phospholipids and phosphonates; whereas the more specialized enzymes are less distributed, such as zinc-β-lactamases, flavodiiron oxidoreductases, and MBLs involved in secondary metabolism, with the important exception of the ubiquitous glyoxalases II. In addition, the structural diversity of membrane-associated MBL proteins has not been explored by experimental structural approaches, despite being involved in fundamental processes like natural transformation and horizontal gene transfer.

**Figure 1.**
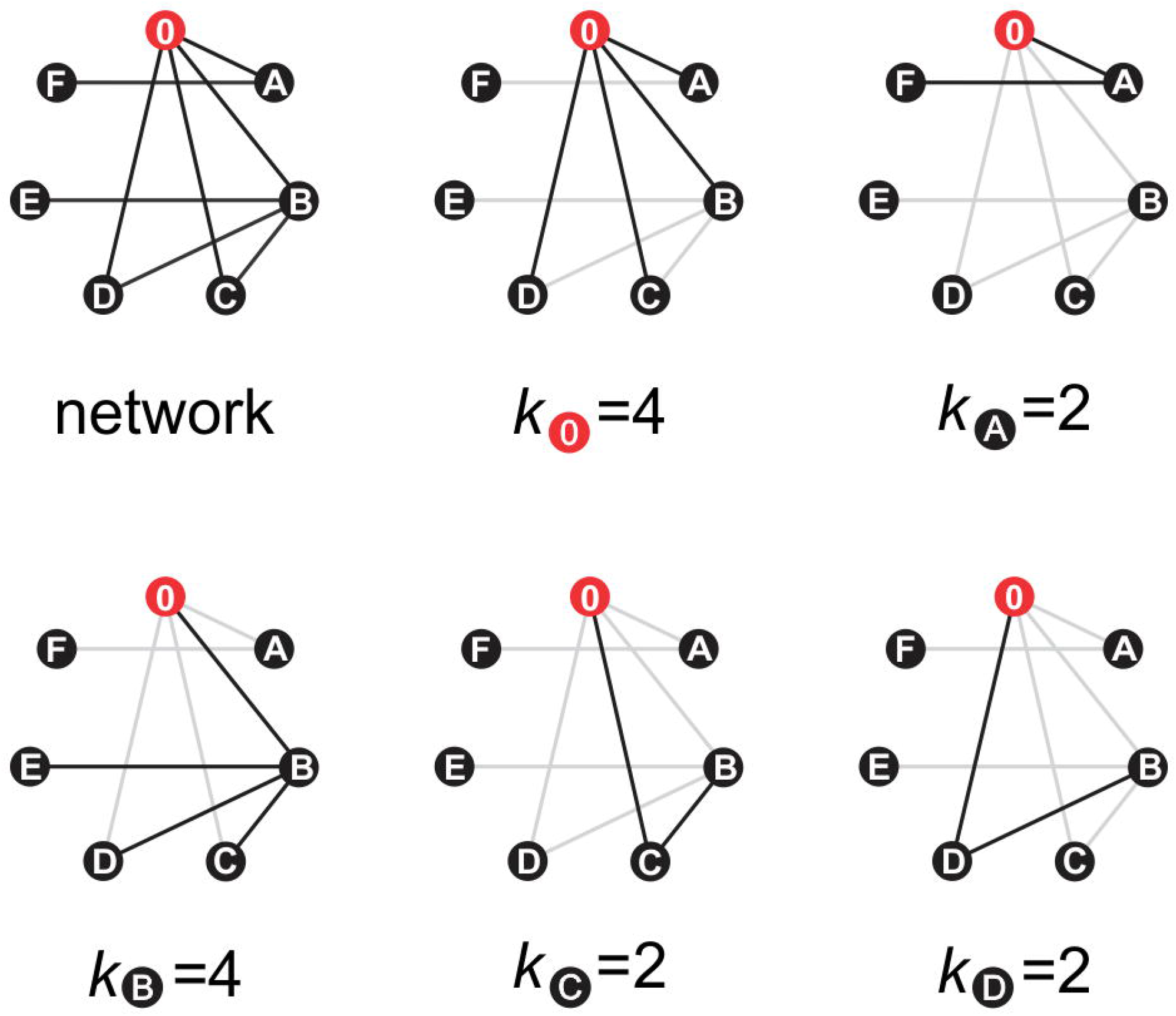
Structural diversity of MBLs. ZBLs like plasmid-borne *Klebsiella pneumoniae* NDM-1 (PDB 4hl2, *left*) comprise a single αββα domain, with the Zn(II) binding site at the bottom of an open groove, accessible to varied β-lactam antibiotics. B1 ZBLs exhibit unusual zinc ligands, including a cysteine residue, which is uncommon in catalytic Zn(II)-binding sites. Instead, habitual MBL metal ligand sets include only histidine and aspartic acid residues. For instance, RNAse J from *Methanolobus psychrophilus* (PDB 6llb, *middle*) comprises a phosphoesterase αββα domain and a β-CASP domain for single-stranded RNA binding. Finally, MBL oxidoreductases such as the flavo-diiron protein ROO from *Desulfovibrio gigas* (PDB 1e5d, *right*) utilize non-heme Fe(II)/Fe(III) for catalysis, exhibiting a more acidic metal ligand set, in combination with an FMN-binding flavodoxin domain, displaying a homodimeric quaternary structure. Metal ions are indicated as numbered spheres. Amino acid side chains follow the coloring scheme of Figure 2.

## MATERIALS AND METHODS

### Structural data harvesting and tanglegram calculation

All MBL protein sequences with available experimentally determined three-dimensional structure were retrieved from the Protein Data Bank (PDB) with the Dali Lite server [12], using structures PDB 2gmn and PDB 3i13 as queries. A set of 105 high-resolution structures was obtained after applying a 90 % sequence similarity cutoff. As well, an unrooted structural dendrogram was obtained for this set with the Dali Lite server all-against-all comparison tool, which calculates a distance matrix of *Z*-scores by aligning the structures all-against-all and outputs a dendrogram derived with the average linkage clustering method [12]. Next, the full amino acid sequence corresponding to each of these 105 structures were retrieved from the UniProt database [13], in order to avoid sequence artifacts like mutations and missing residues often found in PDB files. A structure-guided multiple sequence alignment (MSA) was calculated with Promals3D [14]. This MSA was manually edited with Jalview 2.9 [15] to discard highly gapped regions, by applying a 50 % alignment quality cutoff. The resulting MSA, comprising 105 sequences and 204 columns, was used to calculate a maximum likelihood cladogram with RAxML [16], running at the Cipres server [17]. A best-scoring bootstrapped tree was obtained after 1002 replicates, using the WAG substitution matrix as evolutionary model [18], and was displayed as a consensus cladogram by applying the 50 % majority rule. Finally, in order to compare the consensus sequence-based cladogram with the distance-based dendrogram topologies, a tanglegram matching corresponding taxa was calculated with the Neighbor Net Tanglegram algorithm [19], available in Dendroscope 3.5.9 [20], using the clade of B1&B2 zinc-β-lactamases as outgroup to root each tree. The tanglegram was adapted for display with FigTree 1.4.3 (available at http://tree.bio.ed.ac.uk/software/figtree/) and Corel Draw X7 (Corel). Protein structures were analyzed and graphically represented with PyMOL 1.8 (Schrödinger LLC).

### Sequence data harvesting and SSN calculation

In order to prepare a representative sequence data sample of the MBL superfamily, the PF00753 Pfam database entry was selected as a starting point, which presently comprises 70,367 sequences (release Pfam 32.0, September 2018) [21]. The RP55 representative proteome MSA (62,213 sequences by 1,251 columns) was downloaded and manually edited with Jalview 2.9 [15], by removing truncated and misaligned sequences, highly gapped columns (more than 50 %); and deleting those sequences missing conserved positions corresponding to aspartic acid residues 29, 58, and 134 of human glyoxalase II, which was taken as a reference.

The resulting MSA consisted of 55,076 sequences and 143 columns. Next, the full sequences present in this MSA set were retrieved from the UniProt 2019-10 database [13] and reduced to a final set of 32,418 sequences, by applying a 70 % similarity cutoff with CD-Hit [22] and ensuring that all 105 sequences present in the tanglegram were included. A sequence similarity network (SSN) [23] was then calculated with this 32,418-sequence dataset, using the EFI-EST online tool [24]. The obtained representative node network comprised 15,292 nodes at 40 % sequence similarity, and 762,784 edges at 10^−20^ Blast pairwise similarity threshold. Topology network analysis was performed with NetworkAnalyzer 2.7 [25], as implemented in Cytoscape 3.7.1 [26]. Network statistics plots were prepared with SigmaPlot 12 (Systat Software). All figures were prepared with Corel Draw X7 (Corel).

## RESULTS AND DISCUSSION

### Unearthing ancestral relationships within the MBL superfamily

Tracing the evolutionary history of ancient protein superfamilies is often obscured by the inherent variability of amino acid sequences over long periods. Despite the divergence of primary structure, the three-dimensional fold of polypeptides is less sensitive to mutational events, retaining evolutionary information encoded in the arrangement of secondary structure elements. Thus, experimentally determined structures of proteins offer the possibility of common ancestry inference based on structural homology. Such *phenetic* methods are convenient for comparing proteins with similar folds but highly divergent amino acid sequences, in contrast to MSA-based *cladistic* methods, which are well suited to determine phylogenetic relationships between homologous proteins.

Here, in order to assist the phylogenetic reconstruction of the MBL superfamily, two trees were calculated and compared for a selected set of MBLs with available experimentally determined structures. First, a distance matrix of Dali *Z*-scores comparing all-against-all full-length 105 selected MBL structures was used to construct the corresponding structural phenogram, that is, an unrooted tree whose branch lengths reflect structural similarity relationships between proteins, independently of their amino acid sequence. Next, the amino acid sequences of those 105 polypeptides were retrieved and aligned to construct a maximum-likelihood MSA-based bootstrapped unrooted consensus cladogram, whose topology reflects the sequence homology relationships between extant taxa according to a specific evolutionary model. Both dendrograms were then rooted using the B1&B2 ZBL clade as outgroup, since these enzymes are uniquely divergent MBLs due to their fast-evolving nature. A tanglegram was then calculated with both trees, which consists of a graph of opposing dendrograms with lines connecting equivalent or corresponding taxa, rearranged so that the number of crossing connecting lines is minimal. This type of graph is widely used in Biology to illustrate processes like host-parasite, mutualistic, and symbiotic relationships, where both trees tend to comprise mirror images of each other, as a reflection of their shared topology and evolutionary history. Tanglegrams are used here to explore reciprocal similarities between structure and function of proteins. Since conserved structural features are substantiated by sequence adaptations to perform a specific function, sequence and structure can be assumed to evolve together, and should therefore give rise to dendrograms with the same topology. Crossing connectors between proteins would suggest that conserved residues typical of one group of proteins are found in a scaffold characteristic of different ones. Since the MSA consensus cladogram is not resolved at early nodes, a typical feature of phylogenies of divergent protein families, both trees can be rearranged so that no crossing connecting lines are needed between taxa (Figure 2).

**Figure 2.**
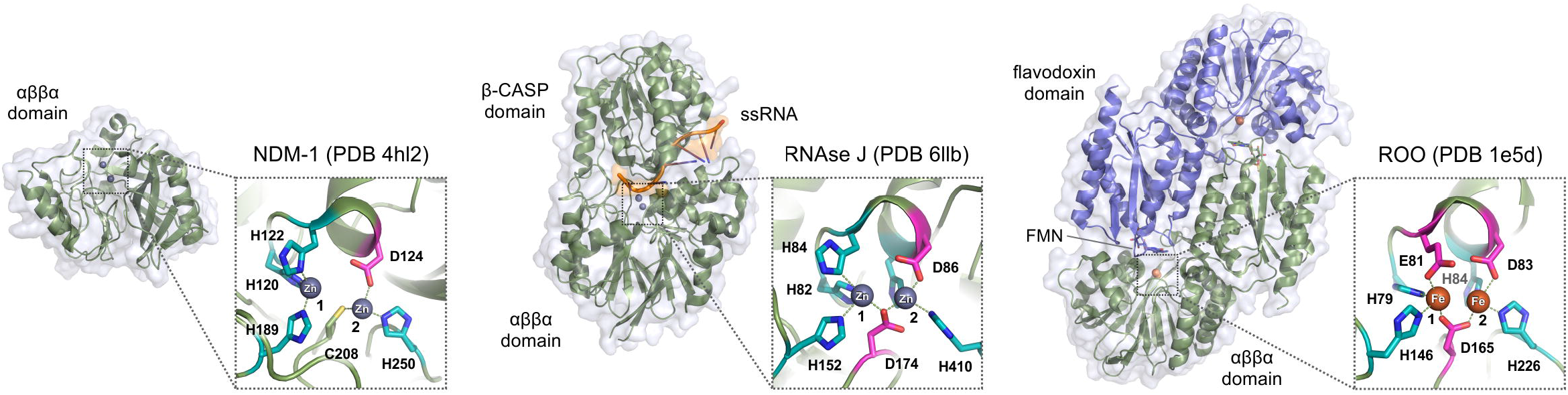
Structure-function tanglegram of the MBL superfamily. Structure-guided phenogram (*left*) and the maximum-likelihood MSA-based bootstrapped consensus cladogram (*right*) of 105 selected MBLs available in the Protein Data Bank. Note that the MSA includes only conserved amino acid residues in the αββα fold, *i.e*. does not take into account additional domains. For each dendrogram, taxa are indicated as representative PDB entries (used for structural phenogram calculation) or UniProt entries (used for MSA and cladogram calculation), respectively. A short version of the MSA is provided, comprising the corresponding sequences sorted with the tanglegram, showing the five MBL fold conserved sequence motifs as histogram logos (*top*), along with short descriptions of common protein names and families (*colored boxes*). While motif 1 contains a conserved aspartic acid residue involved in stabilization of the MBL fold near the active site; motifs 2, 3, 4 and 5 usually contain metal-coordinating residues. In general, Fe(II)/Fe(III) binding sites typical of MBL oxidoreductases exhibit more acidic residues than Zn(II) binding sites, often found in MBL hydrolases. Unusual residues in the MSA are indicated as *red* boxes. Amino acid sequence lengths are variable between these motifs, ranging 6-503 before motif 1 (N-terminus); 9-77 between motifs 1 and 2; 3-23 between motifs 2 and 3; 11-65 between motifs 3 and 4; 14-241 between motifs 4 and 5; and 0-58 after motif 5 (C-terminus). Orange dots in cladogram nodes indicate bootstrap branch support values higher than 50 %.

### Phenetic and cladistic considerations shed light on mutual MBL ancestors

ZBLs comprise a divergent polyphyletic group of MBLs, comprising subclasses B1, B2, and B3 [27]. As shown in the tanglegram and suggested previously [28], B3 ZBLs form a phylogenetically distinct group as compared with B1&B2 enzymes, a clear example of how ZBL activity evolved twice within the superfamily. Motif 2 of B1, B2, and B3 ZBLs are characteristically of the form HxHxDX (where X is not a zinc ligand, typically Arg, Lys or small side chain residues), NxHxDR and HxHxDH, respectively. While B2 ZBLs are typically strict carbapenemases, B1 and B3 ZBLs display low substrate selectivity, and are able to hydrolyze all penicillins, cephalosporins and carbapenems of clinical use. Subclass B1 plasmid-borne ZBLs like IMP-1 (see Figure 2 for UniProt identifiers) became known in the ‘90s for their ability to hydrolyze carbapenems, the latest generation of β-lactam antibiotics available. 30 years later, pathogens expressing B1 enzymes like NDM-1 (Figure 1) still comprise one of the most cumbersome public health issues. In agreement with previous observations, B1&B2 enzymes are closely related and share a recent ancestor, along with a distinctive Zn(II)-binding cysteine at motif 4, supporting antibiotic resistance at limiting Zn(II) concentrations [7]. In contrast, B3 enzymes are typically chromosomal and replace this cysteine with residues unable to coordinate Zn(II) ions, like Ser, Ile, Val, Leu, and Met. In addition, all motif 2 histidines of B3 enzymes become zinc ligands, which is the usual scenario throughout the superfamily. It is worth emphasizing that the HxHxDH motif is the hallmark of the superfamily, and such sequence diversity at motif 2 of ZBLs is rather unusual for a group of enzymes catalyzing the same reaction. This variability likely results from the strong selective pressure exerted by the comparably diverse set of β-lactam antibiotics currently in use.

Notably, the closest structural homologs of B1&B2 ZBLs are not B3 ZBLs but a group of non-heme iron flavoenzymes, known as flavodiiron proteins (FDPs). FDPs like *Desulfovibrio gigas* rubredoxin:oxygen oxidoreductase ROO (Figure 1) [29] comprise a widespread family of prokaryotic oxidoreductases, containing an iron-binding MBL domain and an FMN-binding flavodoxin-like domain [30]. ROO is a terminal reductase, which reduces O_2_ to H_2_O without the risk of producing reactive oxygen species. Other structurally characterized FDPs include *Moorella thermoacetica* and *Escherichia coli* nitric oxide reductases, and the *Giardia intestinalis* oxygen-scavenging enzyme. A typical His-to-Glu mutation appears at motif 2 of FDPs, located at the interface between the isoalloxazine and di-iron moieties, which likely contributes to hold the more acidic Fe(III) species. An unusual metal coordination set is found in *Thermotoga maritima* diiron oxygen sensor ODP [31], where the third histidine of motif 2 is replaced by a glutamine at motif 5. Finally, the divergent class-C type-2 FDPs from *Synechocystis* sp. display mutations at motifs 2, 3 and 4 that prevent binding of any metal ions [32].

The next group comprises phylogenetically distinct proteins structurally related to alkylsulfatases. Type III sulfatases hydrolyze sulfate esters releasing HSO_4_^−^ and the corresponding alcohol. While *Pseudomonas aeruginosa* SdsA1 [33] has preference for primary alcohol sulfates like sodium dodecylsulfate, *Pseudomonas* sp. DSM661 Pisa1 is active on secondary alcohol sulfates, which allowed the discovery that the reaction proceeds with inversion of configuration [34]. Hydrolysis of a secondary alcohol sulfate can proceed through cleavage of C—O or O—S bonds, by nucleophilic attack on the C or S atom, respectively, but only the former can result in inversion of configuration. This is an unprecedented reaction mechanism in the MBL superfamily because the nucleophilic attack occurs on the alcohol carbon by means of an S_N_2 concerted reaction, where HSO_4_^−^ is the leaving group. Thus, MBL *sec*-alkylsulfatases are highly enantioselective enzymes with great potential for application to deracemization processes [35]. In this group, there is also a clade of prokaryotic MBLs of unknown function; the human mitochondrial endoribonuclease LACTB2; and *Pseudomonas* sp. quinolone response protein PqsE. LACTB2 has been shown to use Zn(II) to hydrolyze ssRNA [36]; likely involved in RNA processing specific to mitochondrial function due to its localization and structural homology with bacterial enzymes. PqsE has been shown to bind Fe(II)/Fe(III) *in vitro* and display thiolesterase activity against a CoA-linked intermediate in the biosynthetic pathway of quinolone quorum sensing molecules, although it also contributes to the regulation of bacterial virulence through an unknown mechanism, unrelated to its thiolesterase function [37].

Glyoxalases II (GlxII) and persulfide dioxygenases (PSDO) share a structurally homologous MBL domain, suggestive of common ancestry. This can also be witnessed in the MSA cladogram, where this group forms a separate clade. Human glyoxalase II was the first prototypical MBL to be characterized through X-ray crystallography, disclosing the typical structural features of MBLs. GlxII are thiolesterases that convert *S*-D-lactoylglutathione into D-lactate and glutathione, as part of a ubiquitous methylglyoxal detoxification pathway [38]. The enzyme contains an αββα domain with a consensus HxHxDH motif for binding of two metal ions, reportedly Zn(II) or Mn(II), with an aspartic acid bridge in between. An additional C-terminal domain enables the enzyme to recognize and orient the glutathione moiety for proper hydrolysis, which takes place in the MBL domain metal-binding site. PSDOs are also named ETHE after the human ethylmalonic encephalopathy, a disease that has been linked to mutant PSDO enzymes [39]. Strikingly, while GlxII enzymes harbor a conventional MBL bimetallic center, PSDO enzymes have a single Fe(III) ion at site 1, even though all anticipated metal binding motif residues are conserved. Nevertheless, both enzyme groups catalyze reactions involving glutathione derivatives, *e.g*. 2-hydroxyacyl-glutathione for GlxII and glutathione-persulfide (GSS^−^) for PSDOs, which detoxify sulfide by oxidation to sulfite using molecular oxygen [40]. Some PSDO enzymes like the *Burkholderia phytofirmans* enzyme are fused to rhodanese domains, working instead in sulfur assimilation pathways [41].

The next group comprises at least three phylogenetically distinct structural homologs: quorum-quenching lactonases (QQL), organophosphorus hydrolases (OPH), and human MBLAC1 endonuclease. A number of phenotypes exhibited by bacterial communities are regulated by freely diffusing small molecules signaling cell density. This quorum sensing mechanism is turned off by QQL enzymes like *Bacillus thuringensis* AiiA and *Agrobacterium* sp. AiiB, acting on *N*-acylhomoserine lactones; *Mesorhizobium japonicum* lactonase acting on 4-pyridoxolactone (an intermediate of vitamin B_6_ catabolism); and *Chriseobacterium* sp. AidC lactonase. OPH enzymes like *Pseudomonas* sp. OPHC2 and methylparathion hydrolase MPH are related to QQLs but evolved to hydrolyze phosphoester bonds habitually present in organophosphorus pesticides. Indeed, OPHs may have evolved from QQLs as a resistance mechanism due to the strong selective pressure of these pesticides, resembling how ZBLs evolved to hydrolyze β-lactam antibiotics. Finally, MBLAC1 is a metazoan 3’-end mRNA processing enzyme, acting on stem-loop structures present in histone coding mRNAs [42], constituting the first of many examples of MBL nucleases.

Phosphoesterases comprise the most widespread functional group of the MBL superfamily, hydrolyzing varied phosphoesters like nucleic acids and nucleotides, phosphonates, and phospholipids. Nucleic acid processing enzymes are usually binuclear Zn(II)-dependent hydrolases, such as RNAse J, tRNAse Z, cleavage and polyadenylation specificity factors (CPSF); and DNA repair enzymes like Apollo 5’-exonuclease. These enzymes typically comprise additional domains in a modular fashion that assist the αββα hydrolytic domain at accommodating such large substrates, for instance, the tRNAse Z exosite for tRNA binding [43], β-CASP domains for ssRNA binding [44] (Figure 1), and KH domains for RNA/DNA binding [45]. These modular domains can be either N-terminal, C-terminal, or inserted within the MBL fold. Indeed, the β-CASP domain sequence inserts in the loop holding the conserved His at motif 5, shifting this amino acid about 215 residues towards the C-terminus, making it difficult to find through conventional sequence alignments (*e.g. T. thermophilus* RNAse J). Analogously, the exosite insertion in tRNAse Z shifts the His at motif 5 about 75 residues to the C-terminus (*e.g. E. coli* ZipD). The yeast Trz1 tRNAse Z is an interesting example of a protein with two MBL domains where one of them evolved to improve substrate binding while losing the metal-binding and hydrolytic ability [46] (note that only the catalytic domain of Trz1 was considered in the alignment of Figure 2).

Structurally characterized phosphoesterases devoid of nuclease activity include diverse enzymes like *S. pneumoniae* modular phosphorylcholine esterase CbpE; human *N*-acyl phosphatidyl ethanolamine phospholipase D, NAPE-PLD (the only structurally characterized MBL phospholipase), and dimanganese phosphonatase PhnP from *E. coli*, part of the phosphorus scavenging CP-lyase pathway. Note that PhnP are structurally and phylogenetically related to tRNAse Z enzymes, despite their radically different functions.

*Streptococcus pneumoniae* phosphorylcholinesterase CbpE is localized in the pneumococcal cell envelope [47], and catalyzes the removal the phosphorylcholine from teichoic acids, key components for cell recognition and invasiveness. The divergent *E. coli* manganese-dependent UlaG L-ascorbate-6-P lactonase clusters among phosphoesterases, and has indeed been shown to hydrolyze cyclic nucleotides [48].

Some divergent iron-dependent oxidoreductases cluster at the end of the tanglegram, including *Thermoanaerobacter tengcongensis (C. subterraneus)* Tflp, and *Streptomyces venezuelae* CmlA β-hydroxylase. Tflp contains two Cys residues in the vicinity of the di-iron center, with an Asp-to-Cys mutation at motif 4 (seen so far only in modern B1&B2 zinc-β-lactamases), plus a unique Cys residue following the His residue at motif 5. Complementary spectroscopic assays indicate that Tflp holds an [Fe-S] center under reducing conditions, and structure PDB 2p4z corresponds to an oxidized inactive form. On the other hand, CmlA is a rare β-hydroxylase clustering among phosphoesterases, which hydroxylates L-*p*-aminophenylalanine, a biosynthetic precursor of chloramphenicol.

### SSN analysis suggests that numerous MBL families remain to be characterized

Sequence similarity networks (SSN) have been applied over the past decade to analyze a number of protein superfamilies [23, 49]. Herein, an SSN was calculated for the MBL superfamily using the EFI-EST webserver [24], as described in the Methods section; results are shown in Figure 3. SSNs are graphs with nodes representing protein sequences and edges connecting them, indicating a pairwise sequence similarity at a specified cutoff value. The metric for node similarity calculation at EFI-EST is the Blast *E*-value, which was set to −log(*E*-value) = 20. Unless otherwise stated, nodes are specifically representative nodes, which group several UniProt entries with a 40 % or higher sequence similarity, so that the SSN has fewer edges and is simpler to display graphically. By inspecting the distribution of functionally characterized proteins throughout the SSN it is evident that many MBL families remain to be characterized. In fact, one of the largest clusters in the network comprises proteins involved in DNA internalization and natural competence such as ComEC, for which no structural information is yet available and only one SwissProt entry (*Bacillus subtilis* P39695). The size of connected components (CC) in the SSN follows a power law distribution, with a few clusters encompassing most nodes, and a long tail of many CCs with one or two nodes (Figure 4 A). The largest CC (7259 nodes) includes glyoxalases II, PSDOs, OPHs, QQLs and B3 ZBLs; the second (1962 nodes) includes B1&B2 ZBLs and *sec*-alkyl sulfatases; and the third (1503 nodes) DNA internalization/ComEC proteins; whereas CPSF/β-CASP, tRNAse Z, RNAse J, and FDPs cluster into separate CCs of 673, 350, 333 and 297 nodes, respectively. The remaining 2915 nodes (19 %) include relatively few known MBLs sparsely scattered over 1353 smaller CCs. Analogously, the node degree shows a sharply decaying distribution, skewed towards lowly connected nodes (Figure 4 B). This is probably true for all SSNs for a given alignment score cutoff, since new nodes (proteins) likely become part of existing connected components (families) instead of giving rise to new ones. Nevertheless, the curve is convex up in log-log scale (*inset*), *i.e*. it is not a power law distribution. Only 148 nodes have SwissProt descriptions and 91 nodes have at least one PDB experimentally determined structure (41 nodes have both). As depicted in Figure 3, the majority of nodes with SwissProt and PDB entries describe glyoxalases II, ribonucleases, FDPs, and ZBLs, accounting for 198 out of 15,292 nodes (1.3 %). In other words, 98.7 % of the SSN nodes need experimentally obtained functional and/or structural information so that an accurate annotation can be specified. Given the fast pace at which sequence databases grow, misannotation of macromolecular sequences is an increasingly cumbersome problem [50–52], and relying on entry annotations to define protein families is not a judicious approach.

**Figure 3.**
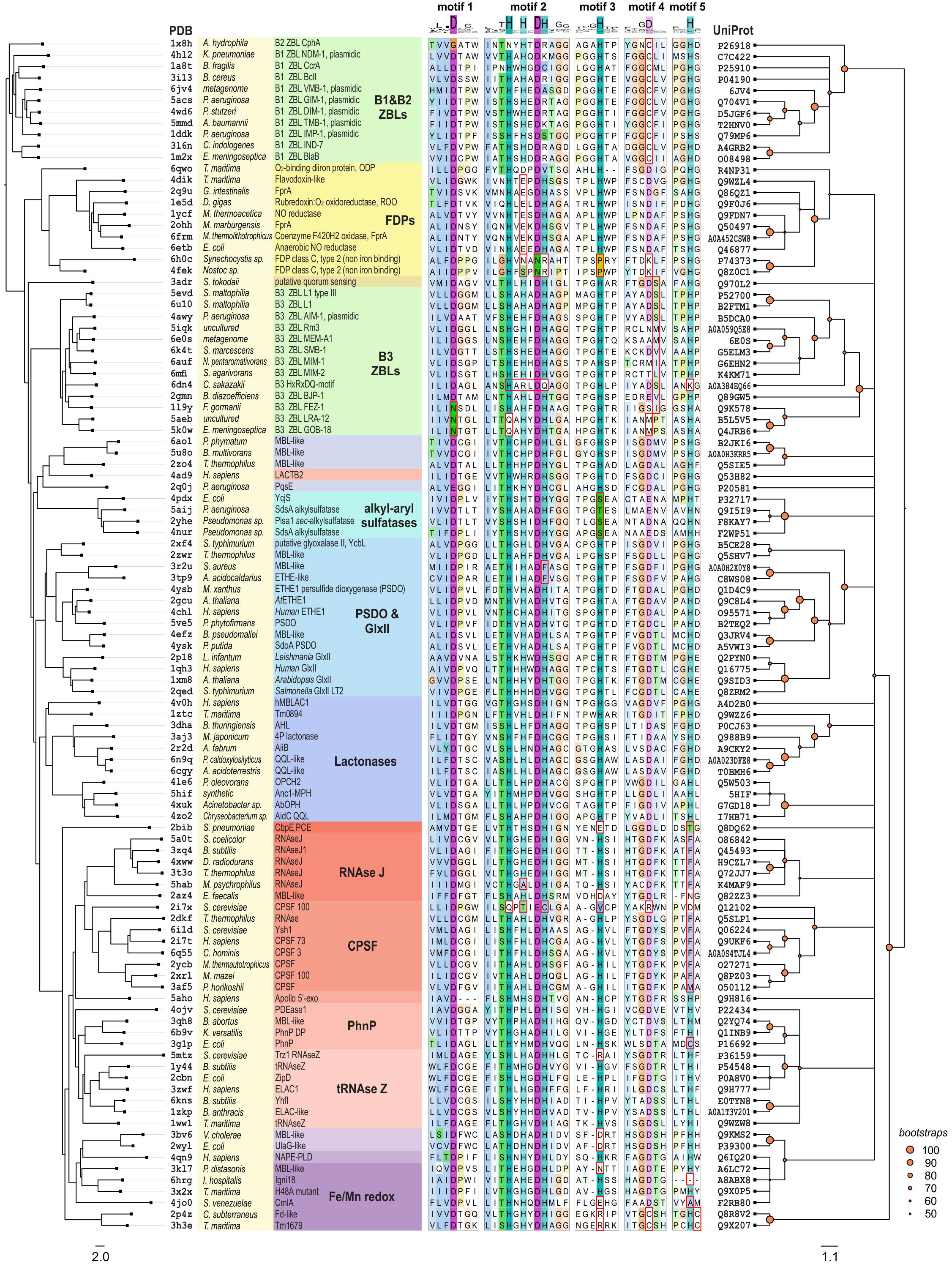
Sequence similarity network (SSN) for representative MBL αββα domains in the Pfam database. The network comprises the amino acid sequence of 32,418 MBLs, expressed as 15,292 representative nodes, grouping connected nodes sharing at least 40 % sequence similarity (each representative node size is scaled by the number of proteins included). Edges between pairs of representative nodes indicate a Blast −log(*E-*value) of 20 or better, which corresponds to a sequence identity of at least ~ 30 %. Note that only MBL αββα domains were considered for Blast score calculations. For comparison, proteins and families included in the tanglegram are indicated. Square nodes indicate sequences with SwissProt and/or PDB descriptions (*inset*). Note that many structurally and functionally characterized proteins do not cluster with the major components of the SSN but are located in isolated components (*bottom*), since their sequence similarity with proteins in major components is on average lower than 30 %. Nodes are organized with the Cytoscape Prefuse Force Directed Open CL layout, and colored by neighborhood connectivity (*top right*).

**Figure 4.**
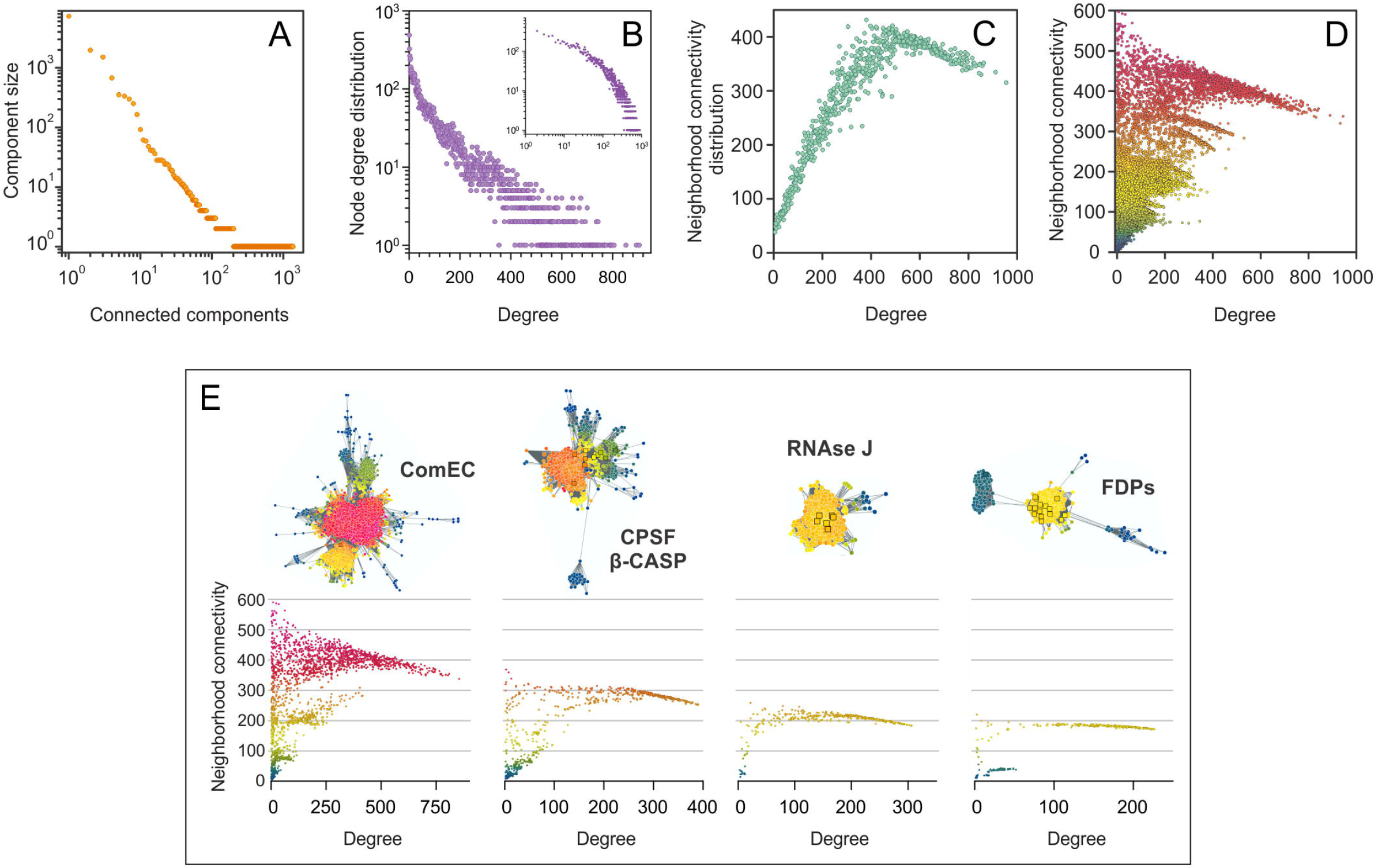
Topological parameters of the MBL superfamily SSN. (**A**) Connected components (CC) are sets of nodes connected by paths of edges. Although a full SSN comprises a single CC, setting an alignment score cutoff leads to a disconnected network aiming to isolate individual protein families, and thereby a set of CCs. The distribution of CC sizes follows a power law, *i.e*. a straight line with negative slope in log-log scale. (**B**) Many natural networks follow a power law distribution of node degrees. However, the SSN node degree distribution is convex up and skewed toward highly connected nodes or nodes with relatively large neighborhoods (Box 1). (**C**) If all NC values are averaged for each degree value, the NC distribution is obtained. A maximum neighborhood connectivity of ~ 400 is observed for *k* ~ 500, which means that, on average, neighborhoods larger or smaller than ~ 500 neighbors are less interconnected. (**D**) Plotting all NC values for each node degree results in a scatter plot with “spikes” for highly interconnected clusters, *i.e*. highly similar groups of proteins (compare with Figure 3, the same coloring was used here). (**E**) Plots of individual NC *vs*. node degree for individual CCs provide a clearer picture of how NC values show an almost inverse linear relationship with connectivity, skewed to larger connectivity values.

### Neighborhood connectivity distribution correlates with protein family clustering

The neighborhood connectivity (NC) statistic was introduced in 2002 by Maslov & Sneppen to describe how sets of highly connected regulatory genes control the expression of lowly connected genes [53] (Box 1). In SSNs, highly interconnected clusters share sequence and, presumably, functional similarity. Thus, members of protein families should have similar connectivities, and coloring nodes by NC provides an intuitive way of visually spotting protein families within CCs. Highly interconnected clusters indicate conserved, highly similar sequences; whereas lowly connected nodes point to rare sequences, proteins underrepresented in the SSN, or simply noise (*e.g*. truncated or incomplete sequences). For a given set of protein sequences, the SSN topology often matches the corresponding phylogenetic tree topology [23]; however, such agreement depends critically on the metrics used for network, MSA, and tree calculation [54]. This is particularly important when comparing divergent sequences sharing few conserved motifs, like the MBL superfamily. For instance, while functional families cluster into distinct clades in the tanglegram, the SSN largest connected component includes most lactonases, glyoxalases II, PSDOs, and B3 ZBLs; and separate clusters are observed for tRNAse Z, RNAse J, and CPSF phosphoesterases (Figure 3). Besides, while B1&B2 ZBLs cluster with alkylsulfatases in the SSN, the tanglegram shows that FDPs are their closest structural homologs. These apparent discrepancies likely reflect the different calculation metrics, *i.e*. sequence similarity for the SSN as opposed to structural homology for the tanglegram. The NC distribution reaches a maximum of ~ 400 for nodes with ~ 500 neighbors (Figure 4C), decaying almost linearly for higher connectivities. Apparently, once clusters reach a maximal connectivity or edges per node, they grow upon addition of new nodes but fewer connections are introduced. This reciprocal linear relationship observed for the full network seems to hold true also for individual clusters: plotting individual NC values reveals linear segments for each cluster, provided that enough nodes are present (Figures 4D&E).

#### Box 1. Neighborhood connectivity.

Unlike many so-called “biological networks” such as protein-protein interaction networks or metabolic networks, SSNs are undirected and do not display self-edges. Then, the *neighborhood* of a node *n* is the set of nodes sharing an edge with *n*; and its *connectivity*, *k*_n_, is the size of its neighborhood, *i.e*. the number of neighbors of *n*. The *degree* of node *n* is the number of edges reaching *n*, which is equivalent to *k*_n_ for SSNs. Then, the *neighborhood connectivity* (NC) of *n*, is defined as the average connectivity of its neighborhood, NC_n_ = Σ(*k*_i_)/*k*_n_ [25, 53]. For example, for a given node 0 (*red*) in the network {0, A, B, C, D, E, F} (*top left*), the neighborhood of 0 is {A, B, C, D} of size *k*_0_ = 4, and the connectivities of each of its neighbors are *k*_A_ = 2, *k*_B_ = 4, *k*_C_ = 2, and *k*_D_ = 2. Then, the neighborhood connectivity of 0 is NC_0_ = (*k*_A_ + *k*_B_ + *k*_C_ + *k*_D_)/*k*_0_ = 2.5. Note that even though nodes E and F are not neighbors of 0, they still influence its NC value by increasing *k*_A_ and *k*_B_. Since members of a protein family are expected to cluster together sharing edges with each other, their neighborhood connectivities will exhibit comparable values. This can be readily appreciated in Figure 3 by coloring nodes according to their NC values. If *N* nodes in an isolated cluster are connected all-to-all, for each node *k* = *N* − 1 (neighbors or edges), all nodes will have a neighbor connectivity NC = *N* – 1. For example, if the network {0, A, B, C, D, E, F} had edges connecting all-to-all its *N* = 7 nodes, each node would have *k* = NC = 6 neighbors (~ *N* for large clusters). In other words, for highly interconnected clusters, the neighbor connectivity approaches to its maximum value, which is roughly the size of the cluster (*cf*. Figure 4C).

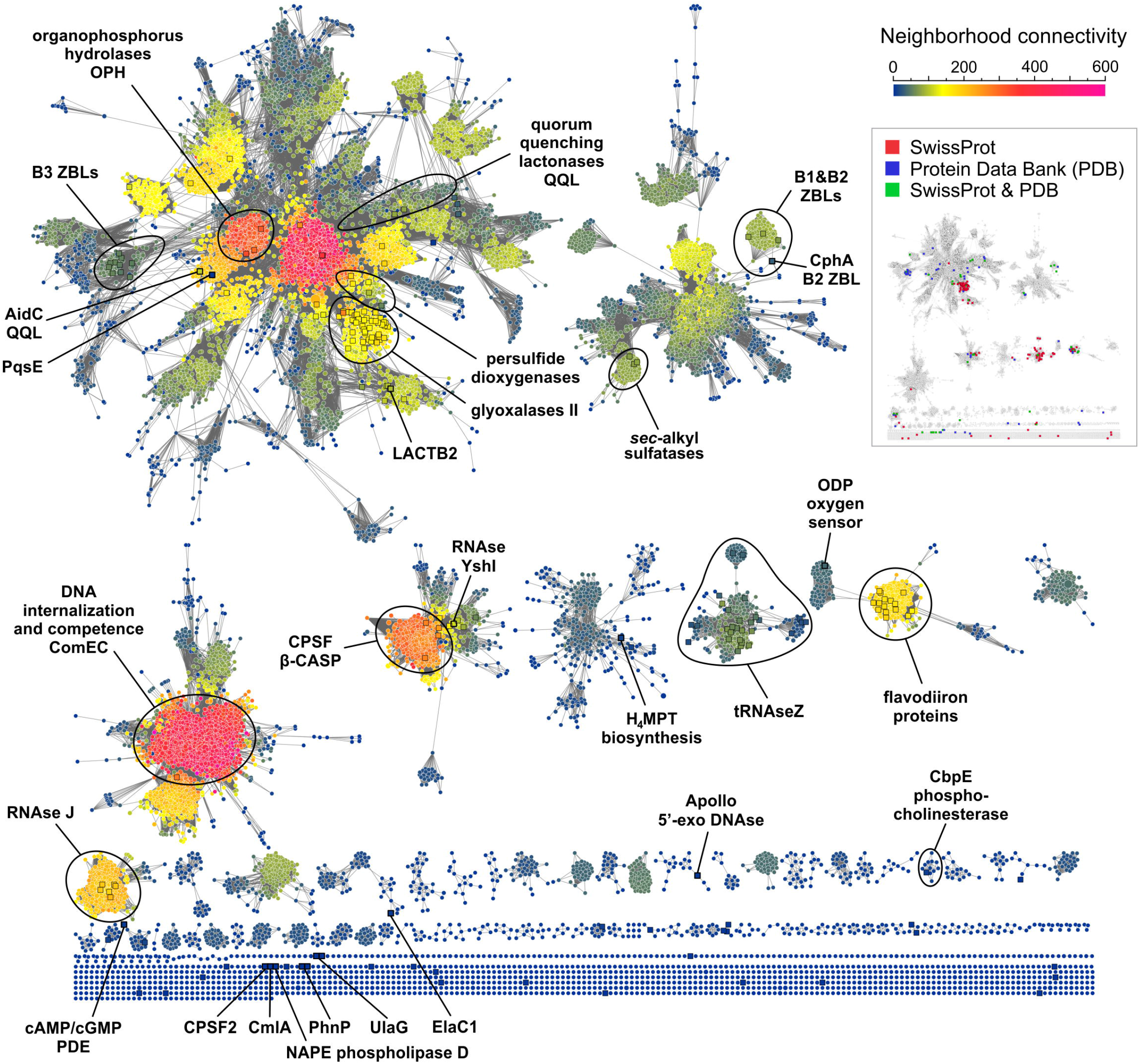

### Concluding remarks

Herein, structural homology and sequence similarity network analysis are used to assist the phylogenetic reconstruction of the MBL superfamily, harnessing the protein three-dimensional arrangement of secondary structure elements as a metric for common ancestry inference. The introduced tanglegram graph disclosed structure and sequence similarity relationships between seemingly unrelated enzymes, which is suggestive of a mutual evolutionary history. In addition, tanglegrams comprise a practical framework for protein structure-function analysis, applicable to study other protein superfamilies as well. Analogously, neighborhood connectivity analysis provides an intuitive picture of the distribution of protein families within the MBL superfamily, suggesting that numerous MBL families remain to be characterized. Indeed, manually annotated entries for proteins with available experimental evidence account for only 1.3 % of the superfamily, underscoring an unfortunately frequent bias of research towards relatively few MBLs. Automated annotation algorithms would benefit from further research on protein sequence similarity networks; establishing their topological features will give rise to improved metrics for protein function estimation.

## Abbreviations

MBL: metallo-β-lactamase
MSA: multiple sequence alignment
SSN: sequence similarity network
PSDO: persulfide dioxygenase
FDP: flavo diiron protein
QQL: quorum quenching lactonase
OPH: organophosphorus hydrolase
ZBL: zinc-β-lactamase
CPSF: cleavage and polyadenylation specificity factor
CC: connected component
NC: neighborhood connectivity

## Acknowledgements

Dr. Liisa Holm is acknowledged for her valuable help with the Dali Lite server. J. M. G. is a staff member of Consejo Nacional de Investigaciones Científicas y Técnicas (CONICET).

